# Compartmentation of photosynthesis gene expression between mesophyll and bundle sheath cells of C_4_ maize is dependent on time of day

**DOI:** 10.1101/2023.04.21.537465

**Authors:** AR Borba, I Reyna-Llorens, PJ Dickinson, G Steed, P Gouveia, AM Górska, C Gomes, J Kromdijk, AAR Webb, NJM Saibo, JM Hibberd

**Affiliations:** Department of Plant Sciences, Downing Street, University of Cambridge, Cambridge CB2 3EA, UK; Instituto de Tecnologia Química e Biológica António Xavier, Universidade Nova de Lisboa, 2780-157, Oeiras, Portugal; Instituto de Biologia Experimental e Tecnológica, 2780-157, Oeiras, Portugal

**Keywords:** C_4_ photosynthesis, mesophyll, bundle sheath, gene expression, maize

## Abstract

Compared with the ancestral C_3_ state, C_4_ photosynthesis enables higher rates of photosynthesis as well as improved water and nitrogen use efficiencies. In both C_3_ and C_4_ plants rates of photosynthesis increase with light intensity and so are maximal around midday. We report that in the absence of light or temperature fluctuations, photosynthesis in maize peaks in the middle of the subjective photoperiod. To investigate molecular processes associated with these changes, we undertook RNA-sequencing of maize mesophyll and bundle sheath strands over a 24-hour time-course. Cell-preferential expression of C_4_ cycle genes was strongest between six and ten hours after dawn when rates of photosynthesis were highest. For the bundle sheath, DNA motif enrichment and gene co-expression analyses suggested members of the DOF and MADS-domain transcription factor families mediate diurnal fluctuations in C_4_ gene expression, and *trans*-activation assays *in planta* confirmed their ability to activate promoter fragments from bundle sheath expressed genes. The work thus identifies transcriptional regulators as well as peaks in cell-specific C_4_ gene expression coincident with maximum rates of photosynthesis in the maize leaf at midday.

## Introduction

In hot and dry environments, C_4_ species can maintain higher rates of photosynthesis and operate higher water and nitrogen use efficiencies than plants that use the ancestral C_3_ cycle (Ghannoum et al., 2010). In C_3_ species the inability of Ribulose 1,5-Bisphosphate Carboxylase/Oxygenase (RuBisCO) to completely distinguish between carbon dioxide (CO_2_) and oxygen (O_2_) leads to competing carboxylation and oxygenation reactions. As temperatures increase and water availability is reduced, the oxygenation activity of RuBisCO becomes more prevalent and so compromises photosynthetic efficiency (Lorimer, 1981; Sedelnikova et al., 2018). More than 60 lineages of land plants have convergently evolved C_4_ photosynthesis and despite some variation in how they concentrate CO_2_ in the leaf, in all cases the likelihood of O_2_ reacting with RuBisCO at the active site of the enzyme is reduced and carbon and energy losses associated with photorespiration suppressed (Bowes et al., 1971; Hatch, 1987; Sage, 2004).

Most C_4_ leaves possess Kranz anatomy, which consists of extensive vascularization combined with an inner wreath of bundle sheath cells and an outer ring of mesophyll cells (Haberlandt, 1904; Langdale, 2011). In C_4_ plants with this leaf anatomy, photosynthetic reactions are normally partitioned between mesophyll and bundle sheath cells. Atmospheric CO_2_ is first converted to bicarbonate (HCO ^-^) by Carbonic Anhydrase (CA) and then assimilated into a four-carbon acid by the O_2_-insensitive PhosphoenolPyruvate Carboxylase (PEPC) in mesophyll cells. Carbon is then shuttled as four-carbon acids to the bundle sheath cells where CO_2_ is released by a C_4_ acid decarboxylase. Three decarboxylases, NAD-dependent Malic Enzyme (NAD-ME), NADP-dependent Malic Enzyme (NADP-ME) and/or PhosphoenolPyruvate Carboxykinase (PEPCK) are known to operate in C_4_ plants to release CO_2_ for re-assimilation by RuBisCO in the Calvin-Benson-Bassham cycle (Hatch, 1987; Kagawa & Hatch, 1974; Y. Wang et al., 2014). The directional transport of organic acids from mesophyll to bundle sheath combined with bundle sheath-preferential accumulation of RuBisCO in C_4_ plants ensure that RuBisCO operates under high CO_2_ concentrations (Sage et al., 2012).

The recruitment of C_4_ genes from the C_3_ photosynthetic pathway required mechanisms that led to patterns of cell-preferential gene expression but also increased transcript levels (Hibberd & Covshoff, 2010; Langdale & Nelson, 1991). These two traits are likely to have evolved independently as they can be controlled by different cis-elements in the same gene (Akyildiz et al., 2007; Kajala et al., 2012; Marshall et al., 1997; Wiludda et al., 2012). Moreover, cell-preferential accumulation of C_4_ enzymes can be specified at different levels of regulation (Gowik et al., 2004, 2017; Heimann et al., 2013; Williams et al., 2016). For example, epigenetic regulation has been documented in the C_4_ monocotyledon Zea mays (maize) where mesophyll-preferential expression of CA and PEPC seems to be regulated by trimethylation of histone H3K4 at analogous gene positions (Heimann et al., 2013). Transcriptional control is also important in C_4_ dicotyledons such as Flaveria bidentis and Gynandropsis gynandra. For example, in F. bidentis mesophyll-preferential expression of PEPC is transcriptionally controlled by cis-elements known as MEM1 and Mesophyll Enhancing Module 1-like (MEM1-like) respectively (Gowik et al., 2004, 2017), and in G. gynandra bundle sheath-preferential accumulation of NAD-ME1, NAD-ME2 and mitochondrial MDH is controlled by a pair of cis-elements that despite being exonic act transcriptionally (Reyna-Llorens et al., 2018). In G. gynandra post-transcriptional regulation is also important, with for example mesophyll-preferential accumulation of CA and Pyruvate,orthophosphate Dikinase (PPDK) being determined through the Mesophyll Expression Module 2 (MEM2) found in 5’ and 3’ untranslated regions (Williams et al., 2016). There is also evidence that translational regulation is important in maintaining cell-specific accumulation of PEPC in maize mesophyll cells, and of RuBisCO in maize and Amaranth bundle sheath cells (Berry et al., 1986, 1988; Chotewutmontri & Barkan, 2020; Wostrikoff et al., 2012).

Despite progress made in understanding global transcriptomic changes associated with the expression of C_4_ genes between cell-types (Aubry et al., 2016; Chang et al., 2012; John et al., 2014b; Ponnala et al., 2014), across developmental gradients (Aubry et al., 2014; Külahoglu et al., 2014; Kümpers et al., 2017) and in response to light (Hendron & Kelly, 2020) to our knowledge very little is known about the effect of photoperiod on cell-preferential gene expression in the C_4_ leaf. To address this, we grew maize under controlled conditions, measured photosynthesis and performed RNA-sequencing from mesophyll and bundle sheath strands over a 24-hour time-course. Although growth conditions were constant, rates of photosynthesis and cell-preferential expression of C_4_ genes varied during the photoperiod. In fact, the largest differences in C_4_ cycle transcript abundance between mesophyll and bundle sheath cells was detected between six and ten hours after dawn, when rates of C_4_ photosynthesis were highest. By integrating a DNA motif enrichment analysis with a gene co-expression network analysis, we identified transcription factors from DOF (DNA binding with One Finger) and MADS (M for MINICHROMOSOME MAINTENANCE FACTOR 1, A for AGAMOUS, D for DEFICIENS and S for Serum Response Factor) families as candidate regulators of bundle sheath-preferential expression. Trans-activation assays in planta confirmed the ability of these DOF and MADS transcription factors to activate promoter fragments of the bundle sheath preferential NADP-ME and PEPCK maize genes.

## Results

### Rates of photosynthesis fluctuate under constant light and temperature

Photosynthetic parameters of C_4_ maize leaves exposed to constant light and temperature were determined 2, 6, 10 and 14 hours after dawn (Figure 1A-E). F_v_/F_m_ values from dark-adapted leaves (Supplemental Table 1) were consistent with those expected from unstressed leaves (Demmig & Björkman, 1987). Despite light intensity being constant, statistically significant variations in assimilation rate were detected (Figure 1A; Supplemental Table 1) with the highest rates occurring ten hours after dawn. The chlorophyll fluorescence parameters φPSII and F_v_’/F_m_’ that report on the operating efficiency of Photosystem II (PSII) and maximum efficiency of PSII without dark adaptation respectively showed slightly different dynamics with values stabilising from two hours after dawn (Figure 1B and 1C). Coincident with the variation in carbon fixation, stomatal conductance increased from dawn to ten hours (Figure 1D). The relative increase in stomatal conductance exceeded that of net CO_2_ fixation, and as a result the intercellular CO_2_ concentration in the leaf increased consistently over the entire fourteen hours of light (Figure 1E). Overall, these data reveal that without alterations in light intensity, photosynthetic parameters in C_4_ maize fluctuate across the day, with higher CO_2_ assimilation at ten hours after dawn (Figure 1A). The trend of increased CO_2_ assimilation, stomatal conductance and intercellular concentration of carbon dioxide until ten hours after dawn contrasted with φPSII and F_v_’/F_m_’ that peaked after only two hours of light (Figure 1A to 1E). To initiate a molecular investigation of processes associated with these alterations to C_4_ photosynthesis over the photoperiod we assessed genome-wide patterns of transcript abundance in mesophyll and bundle sheath cells over a 24-hour period.

**Figure 1.**
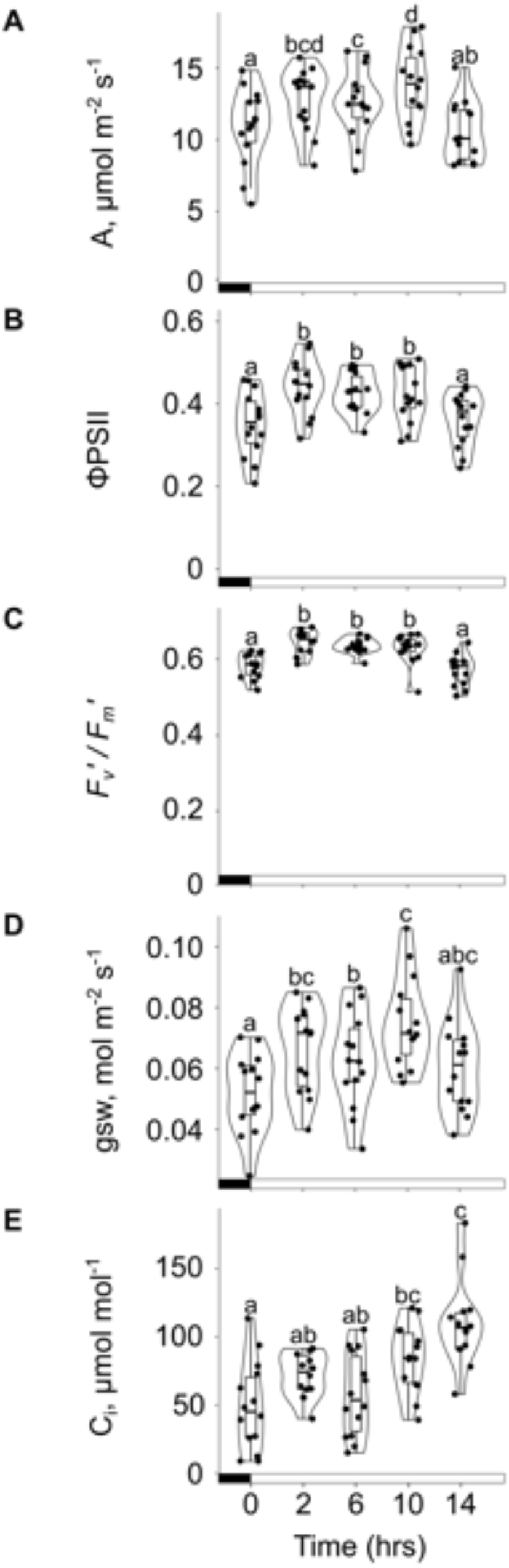
Photosynthetic efficiency in maize fluctuates across the photoperiod. A-E) Violin plots and boxplots showing photosynthetic parameters of light-adapted leaves during constant light and temperature. A) CO_2_ assimilation (A) rate. B) Operating efficiency of Photosystem II (ØPSII). C) Maximum efficiency of PSII photochemistry in the light (F_v_’/F_m_’). D) Stomatal conductance (gsw) to water vapour. E) intercellular CO_2_ concentration (Ci,). Boxplot tails indicate 95% confidence intervals and different letters denote statistically significant differences between time-points determined by One-way repeated measures ANOVA, Tukey test (*p* ≤ 0.05, *n* = 14 biological replicates). Each datapoint represents one biological replicate. Black and white bars in the x-axis denote dark and light periods respectively.

### Compartmentation of C_4_ cycle gene expression varies during the day

RNA was isolated from mesophyll and bundle sheath cells over a 24-hour period and subjected to deep sequencing. Samples were collected at 0, 2, 6, 10, 14, 18 and 22 hours after dawn in a 16 h photoperiod (Figure 2A). 88,521,792 reads were obtained per sample, of which 82% mapped to the maize reference genome B73 AGPv3 (Figure 2B). Quality control for reproducibility showed strong correlation between biological replicates (Pearson’s r > 0.94, Supplemental Figure 1). Principal Component Analysis (PCA) showed that cell-type (mesophyll or bundle sheath) accounted for the first principal component and explained 45% of the variance (Figure 2C). Time of day was associated with the second principal component and accounted for 27% of the variance (Figure 2C). This implies that transcript abundance in the maize leaf is influenced by both cell-type and time of day. To determine whether the spatial patterning of transcripts between mesophyll and bundle sheath cells showed temporal dynamics, differential gene expression analysis was performed at each time-point. The maximum number of differentially expressed genes between these cell-types (12,572) was detected at 6 hours after dawn, whilst the minimum number (9,690) was observed at dawn (0 hrs) (Figure 2D; Supplemental Table 2).

**Figure 2.**
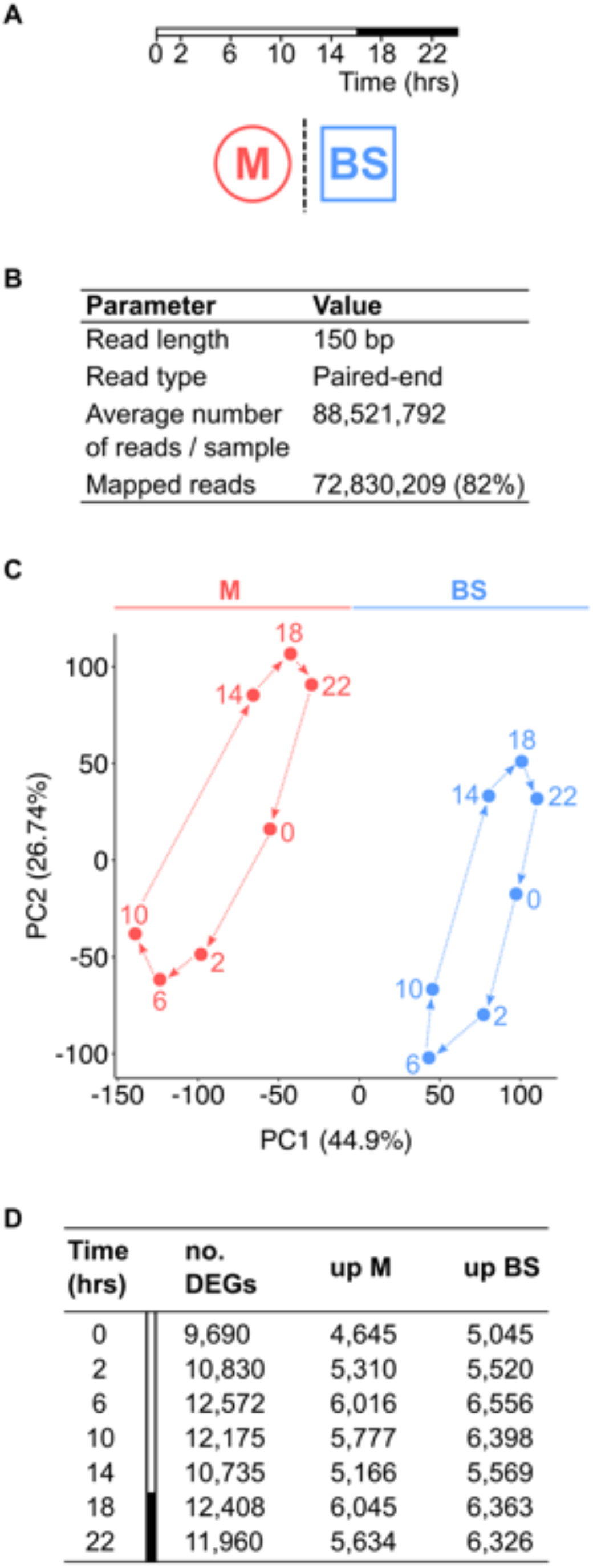
Maize mesophyll and bundle sheath transcriptomes over a diel time-course. A) Mesophyll and bundle sheath transcriptomes were collected over 24-hours. White and black bars denote light and dark periods respectively. B) Transcriptome sequencing parameters. C) Principal Component Analysis of mesophyll and bundle sheath transcriptomes. Principal Component (PC) 1 and PC2 explain 45% and 27% of data variance, respectively. D) Number of differentially expressed genes (DEGs) between mesophyll and bundle sheath cells at each time-point: up-regulated in mesophyll [log_2_(M/BS) > 0] or bundle sheath [log_2_(M/BS) < 0] (DESeq2 differential expression testing with multiple test corrected p-adj < 0.01). M and BS represent mesophyll and bundle sheath cells, respectively.

Core components of the maize circadian oscillator changed over the time-course as would be expected from analysis of C_3_ species. Maize orthologs for circadian oscillator components were defined using OrthoFinder (Emms & Kelly, 2019) using proteomes of Arabidopsis thaliana (C_3_) Zea mays (C_4_), Oryza sativa (C_3_), Triticum aestivum (C_3_), Brachypodium distachyon (C_3_), Setaria italica (C_4_) and Sorghum bicolor (C_4_) as input (Supplemental Figure 2A; Supplemental Table 3). Many circadian oscillator genes in A. thaliana had more than one ortholog in maize (Supplemental Table 3), consistent with the multiple gene duplications in the maize lineage since it diverged from their last common ancestor (Lee et al., 2013). Specifically, Arabidopsis Pseudo-Response Regulator 7 (PRR7, AT5G02810) had three orthologs in maize, hereafter referred to as PRR7.1 (GRMZM2G005732), PRR7.2 (GRMZM2G033962) and PRR7.3 (GRMZM2G095727) (Supplemental Figure 2B). By contrast, Arabidopsis PRR3 (AT5G60100), PRR5 (AT5G24470), PRR9 (AT2G46790) and a CCT motif family protein (AT2G46670) were part of the same clade and shared two orthologs PRR3/5/9.1 (GRMZM2G179024) and PRR3/5/9.2 (GRMZM2G367834) in maize (Supplemental Figure 2C). As expected, maize circadian oscillator genes were expressed in temporal waves with CCA1/LHY.1 and CCA1/LHY.2 transcripts peaking six hours after dawn (Supplemental Figure 2D). The peak in CCA1/LHY transcript abundance was followed by sequential accumulation of PRRs. For example, transcripts of PRR7.1 to PRR7.3 accumulated between six and ten hours of light, and PRR3/5/9.1 and PRR3/5/9.2 peaked at ten and fourteen hours after dawn (Supplemental Figure 2D). A rise in abundance was then observed for the evening/night transcripts LUX ARRHYTHMO (LUX) and TIMING OF CAB EXPRESSION 1 (TOC1.1 to TOC1.6) such that they peaked two hours after the dark period (Supplemental Figure 2D). Despite EARLY FLOWERING 3 (ELF3) being an evening component in A. thaliana (Nusinow et al., 2011) in maize ELF3 transcript abundance was slightly higher during the day (Supplemental Figure 2D). This observation is consistent with previous observations showing ELF3 peaking near dawn in sorghum, foxtail millet, rice and wheat (Zhao et al., 2012; Lai et al., 2020; Wittern et al., 2022). Notably, whilst most core components of the maize circadian oscillator appeared to be partitioned equally between the two cell-types, transcripts for PRR7.1 to PRR7.3 and ELF3 were more abundant in bundle sheath cells across the day (Supplemental Figure 2D).

To investigate whether the circadian clock modulates C_4_ photosynthesis, we measured photosynthetic activity under one light-dark cycle followed by 72 hours of a light regime that consisted of 40 minutes light and 20 minutes darkness (Supplemental Figure 3). Rhythmic oscillations with near 24 h free running circadian periods were detected in the chlorophyll fluorescence parameters F_m_, F_v_/F_m_, φPSII and F_v_’/F_m_’ that report on the maximum yield of fluorescence, maximum quantum efficiency of PSII photochemistry, operating efficiency of PSII and maximum efficiency of PSII (empirical p-value < 0.01, Supplemental Figure 3A-D). φPSII and F_v_’/F_m_’ (Supplemental Figure 3C and 3D) showed similar dynamics to those observed in the dark-light cycle (Figure 1B and 1C) with higher values occurring between two and ten hours after dawn. The 24 h cycles of photosynthetic parameters in these conditions is indicative of circadian regulation. To define groups of genes with maximal transcript abundance at different times of day in each cell-type, k-means clustering was performed (Supplemental Table 4). This identified fifteen clusters of genes that were divided in five groups based on their peak in expression (Figure 3A; Supplemental Table 4). Of the fifteen clusters defined, three of them did not show a strong cell-specific profile (clusters 5, 9 and 11). On the other hand, we observed a clear separation of the clusters defined by the peaks of activity and cell type-preferential expression for the remaining twelve clusters (Figure 3A). To better understand these broad alterations in gene expression, Gene Ontology (GO) enrichment analysis was performed on each cluster (Supplemental Figure 4; Supplemental Table 5). Signalling cascades peaked early in the morning in both cell-types. Later on, transcripts associated with chloroplast organisation, photosynthesis and response to light peaked in mesophyll cells, whilst transport peaked in the bundle sheath. The activation of genes involved in transcription, translation, and protein metabolism was observed during the transition to the dark period (Supplemental Figure 4; Supplemental Table 5).

**Figure 3.**
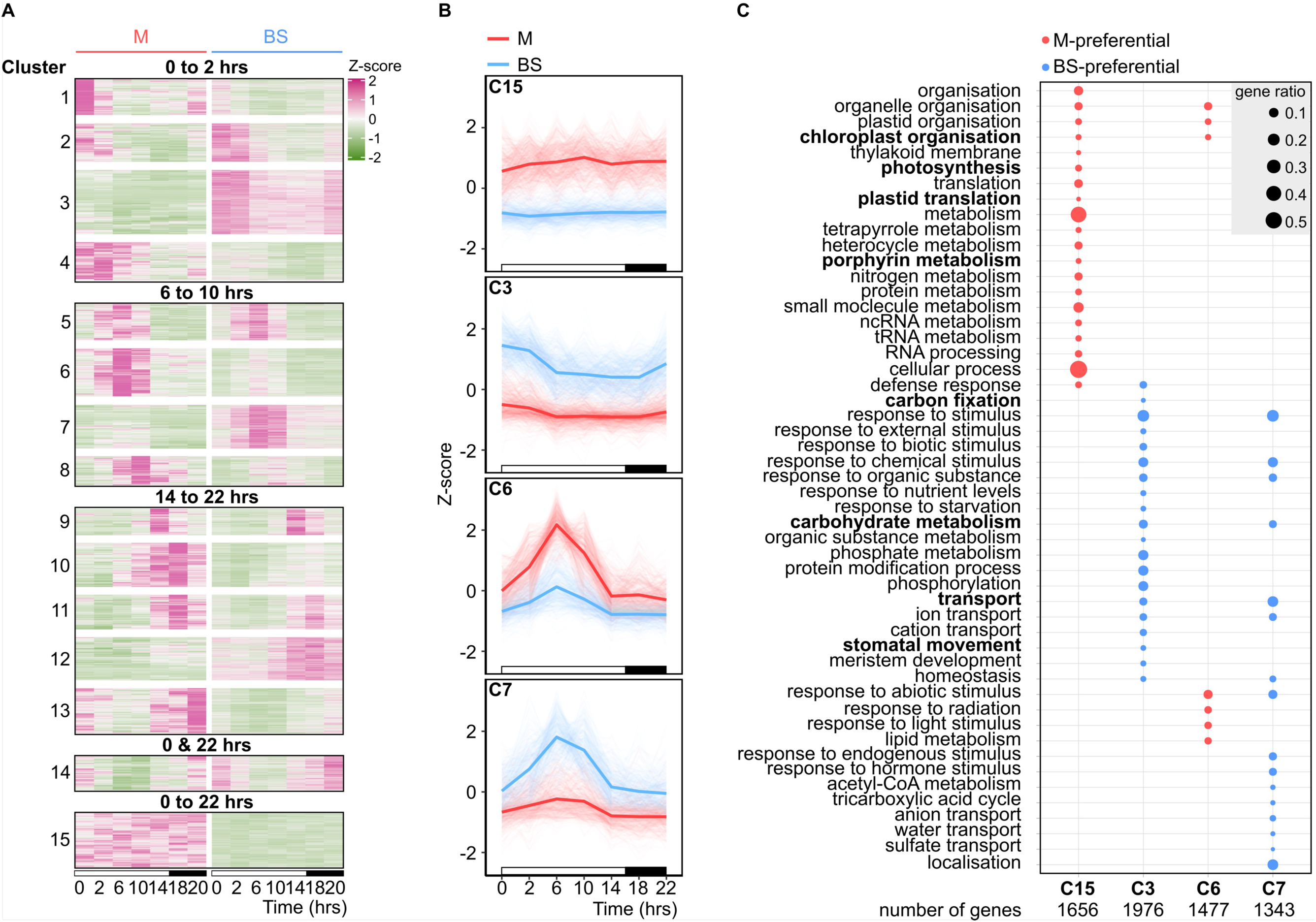
Gene Ontology terms associated with time of day and cell type in the maize leaf. A) Heatmap illustrating profiles of transcript abundance of co-expressed genes in mesophyll and bundle sheath cells across the diel time-course. Clusters are grouped based on the time they peak (from dawn to 2 hours of light, 6 to 10 hrs, 14 to 22 hrs, dawn and 22 hrs, and dawn to 22 hrs). *x*-axis represents time and *y*-axis Z-score. High to low Z-score values are shown as pink to green. B) Line plots representing the diel transcript abundance profile of clusters 15, 3, 6 and 7 in mesophyll and bundle sheath cells across the diel time-course. Thick lines denote the mean of Z-score values in mesophyll or bundle sheath. The *x*-axis represents time-points and the *y*-axis Z-score values. White and black bars in the *x*-axis denote light and dark periods, respectively. C) Dot plot showing the twenty categories of biological processes with highest significance for clusters 15, 3, 6 and 7 (FDR:≤S 0.01). Gene ratio represents the proportion of genes assigned to a functional category in a cluster. M and BS represent mesophyll and bundle sh^1^eath cells, respectively.

Clusters 3, 6, 7 and 15 contained transcripts that showed the most distinct differences in expression between mesophyll and bundle sheath cells (Figure 3B) and so we assessed the nature of genes encoding these transcripts. Cluster 15 contained genes preferentially expressed in the mesophyll throughout the diel time-course and was strongly enriched in biological processes such as chloroplast organisation, photosynthesis, plastid translation and porphyrin metabolism (Figure 3B and 3C; Supplemental Table 5). In contrast, cluster 3 was bundle sheath-preferential and enriched GO terms included carbon fixation, carbohydrate metabolism, transport, and stomatal movement (Figure 3B and 3C; Supplemental Table 5). Interestingly, chloroplast organisation was also enriched in cluster 6 of mesophyll-preferential genes that peaked at six hours after dawn, and cluster 7 that contained genes involved in carbohydrate metabolism and transport that were bundle sheath-preferential (Figure 3B and 3C; Supplemental Table 5).

Consistent with enrichment in the photosynthesis GO term, cluster 15 contained genes from both the core C_4_ and Calvin-Benson-Bassham cycles [PHOSPHOENOLPYRUVATE CARBOXYLASE (PEPC), ASPARTATE AMINOTRANSFERASE from mesophyll (AspAT (M)) and PYRUVATE,ORTHOPHOSPHATE DIKINASE (PPDK) and TRIOSEPHOSPHATE ISOMERASE (TPI)] (Supplemental Table 6). Moreover, cluster 3 was enriched in C_4_-related genes [NADP-DEPENDENT MALIC ENZYME (NADP-ME); RIBULOSE 1,5-BISPHOSPHATE CARBOXYLASE/OXYGENASE ACTIVASE (RCA), FRUCTOSE-1,6-BISPHOSPHATASE (FBP), TRANSKETOLASE (TKL), RIBULOSE-PHOSPHATE3 EPIMERASE (RPE), SEDOHEPTULOSE-1,7-BISPHOSPHATASE (SBP) and PHOSPHORIBULOKINASE (PRK)]. This was also the case for clusters 6 and 7 [with cluster 6 containing CARBONIC ANHYDRASE (CA); GLYCERALDEHYDE 3-PHOSPHATE DEHYDROGENASE B SUBUNIT (GAPDH(B)), and cluster 7 containing PHOSPHOENOLPYRUVATE CARBOXYKINASE (PEPCK); RuBisCO SMALL SUBUNIT-3m (RBCS3m), GLYCERALDEHYDE 3-PHOSPHATE DEHYDROGENASE A SUBUNIT (GAPDH(A)) and FRUCTOSE BISPHOSPHATE ALDOLASE (FBA)] (Supplemental Table 6).

Transcript abundance of C_4_ cycle genes in clusters 3, 6, 7 and 15 varied over the diel time-course and tended to peak during the light period (Figure 4A). Maximal transcript abundance of most C_4_ cycle and also Calvin-Benson-Bassham cycle genes took place between six and ten hours of light (Figure 4A). Indeed, during the first ten hours of light there was a gradual increase in the statistical significance associated with the extent to which C_4_ and Calvin-Benson-Bassham cycle transcript abundance was partitioned between mesophyll and bundle sheath cells (Figure 4B). Taken together these data reveal a striking variation in the extent to which C_4_ photosynthesis genes are preferentially expressed in mesophyll or bundle sheath cells over the day.

**Figure 4.**
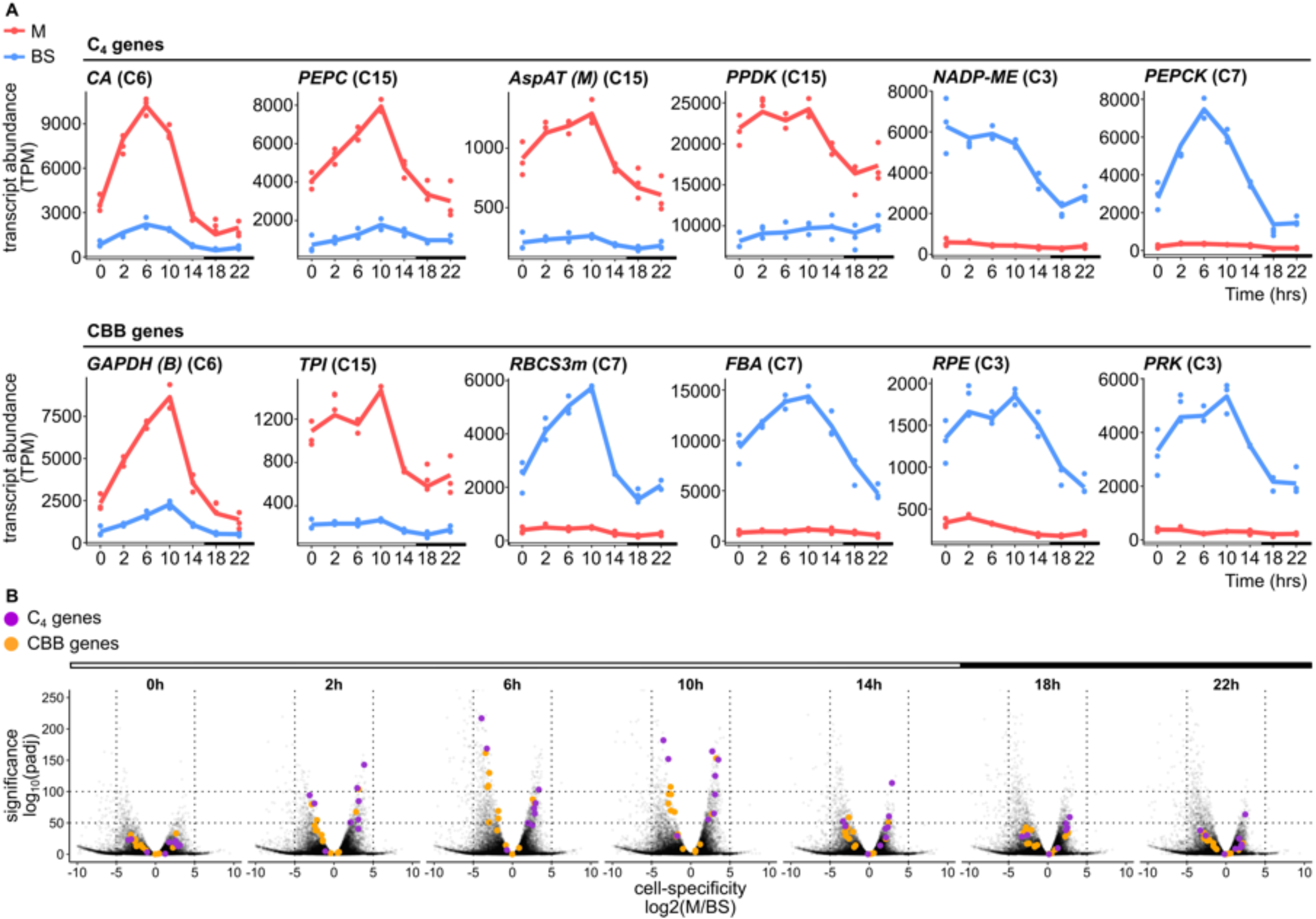
Cell specificity of C_4_ cycle and Calvin-Benson-Bassham cycle transcripts oscillates over the time-course. A) C_4_ genes and Calvin-Benson-Bassham cycle (CBB) genes present in clusters 15, 3, 6 and 7. *x*-axis depicts time and *y*-axis shows transcript abundance in Transcripts Per Million (TPM). White and black bars denote light and dark periods. Gene names are followed by cluster number in parentheses. B) Volcano plots showing the distribution of adjusted *p*-values in relation to the fold-change between mesophyll and bundle sheath cells. Purple and orange circles denote C_4_ and Calvin-Benson-Bassham cycle genes respectively and grey datapoints the remaining transcriptome.

### Members of the DOF and MADS-domain transcription factor families as regulators of bundle sheath-preferential expression of C_4_ and Calvin-Benson-Bassham cycle genes

We next sought to use the RNA-seq time-course to identify cis-elements and trans-factors linked to the control of C_4_ gene expression. Thus, to identify potential regulators in cis and trans of genes in clusters associated with the C_4_ and Calvin-Benson-Bassham cycles (clusters 15, 6, 3 and 7) we performed a motif enrichment analysis using a set of 259 DNA-binding motifs for Z. mays from the PlantTFDB (Jin et al. 2017); Figure 5A, Supplemental Figure 5, Supplemental Table 7). Of the motifs tested less than 10% were enriched in at least one of the four clusters (Fisher’s exact test, p-value < 0.01, Supplemental Figure 5, Supplemental Table 7). Mesophyll-preferential clusters were enriched in only three motifs. Whilst cluster 15 was enriched in the CPP-transcription factor 1 (CPP1) motif, cluster 6 was enriched in G2-like-transcription factor 56 (GLK56) and MYB-transcription factor 138 (MYB138) motifs (Figure 5A; Supplemental Figure 5). The GLK56 transcription factor is a known regulator of the circadian clock (Zhao et al., 2023), activating CCA1 and being co-regulated with TOC1. GLK56 expression peaked at 18 hrs similar to TOC1 orthologs (Supplemental Figures 2D and 5). However, bundle sheath-preferential clusters showed a higher number of enriched motifs. Cluster 3 was enriched in DNA-binding One Zinc Finger 21 (DOF21) and MYB-transcription factor 14 (MYB14) motifs whilst cluster 7 showed enrichment in NLP-transcription factor 13 (NLP13), KNOTTED 1 (KN1), ABI3-VP1-transcription factor 19 (ABI19) and several members of the HSF and SBP transcription factor families (Figure 5A). Moreover, both clusters shared an enrichment for a pair of BBR motifs (BBR3 and BBR4) as well as motifs recognised by DNA-binding One Zinc Finger 2 (DOF2) and MADS-domain protein 1 (MADS1) (Figure 5A; Supplemental Figure 5). This finding suggests that these transcription factors might contribute to bundle sheath-preferential gene expression across the day.

**Figure 5.**
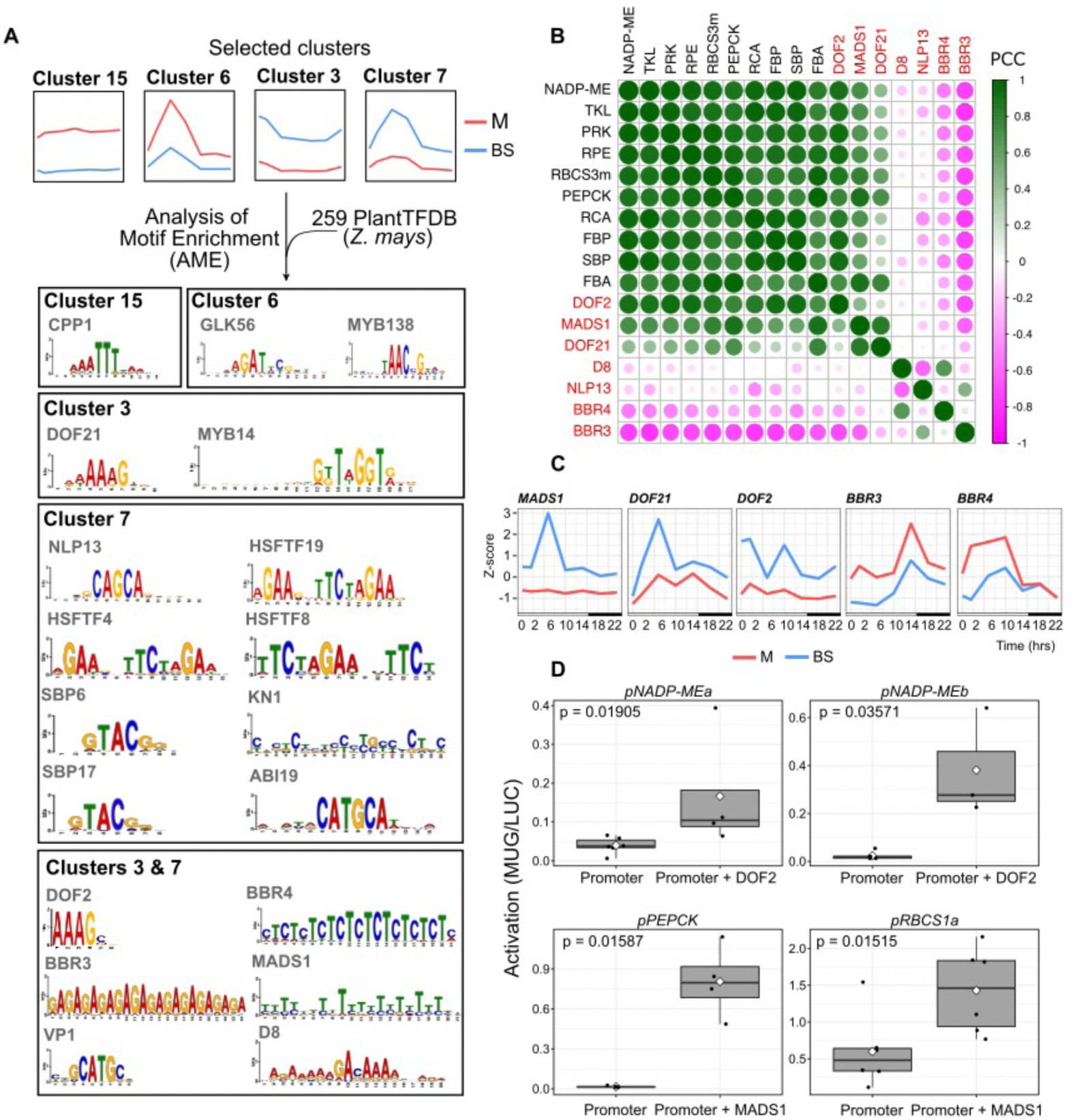
Motifs and transcription factors associated with cell-preferential gene expression. **A)** Four clusters were selected for analysis. DNA-binding motifs enriched in mesophyll clusters 15 and 6, or bundle sheath clusters 3 and 7. **B)** Heatmap illustrating Pearson’s correlation coefficient (PCC) values for bundle sheath-preferential photosynthesis genes in clusters 7 and 3 and candidate transcriptional regulators. DNA-binding One Zinc Finger 2 (DOF2), GRMZM2G009406; MADS-domain protein 1 (MADS1), GRMZM2G171365; DNA-binding One Zinc Finger 21 (DOF21), GRMZM2G162749; Dwarf Plant 8 (D8), GRMZM2G144744; NLP-transcription factor 13 (NLP13), GRMZM2G053298; BBR/BCP-transcription factor 4 (BBR4), GRMZM2G118690; BBR/BCP-transcription factor 3 (BBR3), GRMZM2G164735. **C)** Line plots of diel transcript abundance for candidate regulators of bundle sheath-preferential photosynthesis genes. *x*-axis shows time and *y*-axis Z-score. White and black bars in the *x*-axis denote light and dark periods, respectively. M and BS represent mesophyll and bundle sheath cells. MADS-domain protein 1 (MADS1), GRMZM2G171365; DNA-binding One Zinc Finger 21 (DOF21), GRMZM2G162749; DNA-binding One Zinc Finger 2 (DOF2), GRMZM2G009406; BBR/BCP-transcription factor 3 (BBR3), GRMZM2G164735; BBR/BCP-transcription factor 4 (BBR4), GRMZM2G118690. **D)** Box plots showing promoter activation of bundle sheath-preferential genes *NADP-ME* (cluster 3), *PEPCK* (cluster 7) and *RBCS* (cluster 7) by transcription factors DOF2 and MADS1. Different letters represent statistically significant differences (*P* < 0.05) as determined by two-sided, pairwise t-tests. n=6 for *pNADMEa*, *pRBCS* and *pRBCS*+MADS1, n=5 for *pNADPMEb* and *pPEPCK*, n=4 for *pNADMEa*+DOF2 and *pPEPCK*+MADS1 and n=3 for *pNADMEb*+DOF1.

To further investigate links between enriched motifs and photosynthesis genes present in clusters 15, 6, 3 and 7, a gene co-expression network was built between the corresponding transcription factors and photosynthesis genes containing motif hits (Supplemental Figure 6). Although we started with the four clusters associated with either mesophyll or bundle sheath strands, for two reasons we focussed on those defined by bundle sheath-preferential expression (clusters 3 and 7). First, we did not detect any motif hits for photosynthesis genes present in mesophyll cluster 6. Second, poorly expressed transcription factors (Transcript Per Million reads < 5) were removed and this meant that photosynthesis genes from cluster 15 were also no longer present in the network (Supplemental Figure 6). Pearson’s correlation coefficient was used to define negative or positive co-expression between bundle sheath-preferential photosynthesis genes and candidate transcriptional regulators (Figure 5B). DOF2, MADS1 and DOF21 were positively co-expressed with bundle sheath-preferential photosynthesis genes in cluster 3 (NADP-ME, TKL, PRK, RPE, RCA, FBP, SBP and FBA) and cluster 7 (RBCS3m and PEPCK), whilst BBR4 and BBR3 showed negative co-expression correlation with these photosynthesis genes (Figure 5B, Supplemental Table 7). These relationships are underpinned by MADS1 and DOF21 being preferentially expressed in bundle sheath cells and peaking six hours after dawn, whilst BBR3 and BBR4 peaked towards the end of the light period and were preferentially expressed in mesophyll cells (Figure 5C). We therefore hypothesized that MADS1 and DOF21 act as positive transcriptional regulators of bundle sheath expressed genes whilst BBR3 and BBR4 act to repress these genes in mesophyll cells. To initiate testing, a trans-activation assay in Nicotiana benthamiana was performed. Promoter fragments from the NADP-ME, RBCS and PEPCK genes containing the relevant motifs generated low levels of autoactivation (Figure 5D) and so we were not able to test for negative regulation by BBR3 and BBR4. However, the DOF2 and MADS1 transcription factors activated short promoter fragments of the NADP-ME, PEPCK and RBCS genes containing their cognate motifs (Figure 5D). The combined findings that DOF2 and MADS1 are co-expressed with C_4_ genes, that their DNA binding sites are found in C_4_ promoters, and that they trans-activate expression in planta indicate that these transcription factor families likely play a role in enhancing C_4_ gene expression in the bundle sheath during the day.

## Discussion

### Variation in the rate of C_4_ photosynthesis over the day is influenced by circadian oscillations

Our analysis shows that under moderate illumination and a constant light regime similar to those used in A. thaliana, barley and wheat to study circadian oscillations (Dakhiya et al., 2017; Litthauer et al., 2015, 2016; Wittern et al., 2022) photosynthetic rates vary in maize. These findings are therefore consistent with the fact that photosynthesis in C_3_ species is modulated by the circadian oscillator (Dodd et al., 2005) and our analysis of chlorophyll fluorescence quenching in maize supports this notion. The circadian oscillator also regulates stomatal conductance in C_3_and C_4_ leaves (Resco de Dios & Gessler, 2018). Consistent with the circadian regulation of stomatal conductance and photosynthetic efficiency as has been reported in C_3_ species (Dodd et al., 2005; Harmer et al., 2000) fourteen hours after dawn all photosynthetic parameters except intercellular concentration of carbon dioxide appeared to decline. In this study, CO_2_ assimilation and stomatal conductance followed a different trajectory compared with φPSII and F_v_’/F_m_’, with the former reaching maximum values at ten hours and the latter at two hours after dawn. This apparent increase in rates of CO_2_ assimilation during the day compared with activity of the photosystems, could be because the carbon concentrating mechanism operating in maize is not completely CO_2_-saturated before ten hours. If this is the case, stomatal opening over the day would allow increased intercellular concentration of carbon dioxide and thus higher CO_2_ assimilation. It is also possible that the efficiency of carbon assimilation rises during the day, and stomata respond to this to maintain CO_2_ supply. A third possibility is that at dawn C_4_ photosynthesis operates exclusively with NADP-ME for decarboxylation. As the day progresses, the sustained activity of PSII provides sufficient NADPH in the bundle sheath for PEPCK to act as a second decarboxylase. These hypotheses could be mediated by modifications to the transcriptional activity of genes involved in the C_4_ pathway.

### Compartmentation of C_4_ gene expression between mesophyll and bundle sheath varies over the day

Over the light and dark period we detected statistically significant variance in transcript abundance in mesophyll and bundle sheath strands. Although the main factor explaining this was associated with preferential accumulation of transcripts to either the mesophyll or bundle sheath, time of day also had a significant effect. Thus, although C_4_ cycle transcripts are differentially expressed between the two cell-types (Chang et al., 2012; Li et al., 2010; Tausta et al., 2014), this compartmentation is more dramatic at midday prior to the highest rates of photosynthesis.

Differences in transcript abundance between the two cell-types were associated with the mesophyll being biased towards strong expression of components of the photosynthetic electron transport chain as well as responses to far red, red, and blue light. In contrast, GO terms over-represented in the bundle sheath were involved in carbon fixation and transport. These findings are consistent with the fact that maize mesophyll cells contain both Photosystems I and II whilst bundle sheath strands contain RuBisCO and fail to accumulate significant amounts of Photosystem II (Meierhoff & Westhoff, 1993). Not only did transcripts encoding components of the core photosynthetic apparatus vary in the extent to which they were compartmented between mesophyll and bundle sheath cells, but this was also the case for transcripts associated with signal transduction pathways and stomatal movement. In both cases their transcripts tended to peak prior to those associated with carbon fixation.

### The role of MADS-domain and DOF transcription factors in activation of C_4_ genes in the bundle sheath

In addition to biological processes being enriched in either mesophyll or bundle sheath strands and the extent of this being time of day-dependent, we observed spatiotemporal changes to transcripts encoding multiple transcription factor families. To better understand how transcriptional regulators control the expression of C_4_ and Calvin-Benson-Bassham cycle genes, we performed a motif enrichment analysis on photosynthesis genes followed by a gene co-expression analysis between photosynthesis genes that showed enrichment in DNA-binding motifs and their target transcription factors. This predicted that shared cis-elements and trans-factors control bundle sheath-specificity of genes from both the C_4_ and Calvin-Benson-Bassham cycles, which might ensure spatial and temporal coordination between these two photosynthetic cycles. Although to our knowledge the specific cis-elements and transcription factors identified here have not previously been implicated in controlling C_4_ photosynthesis, there are several reports showing that multiple C_4_ genes can be regulated by the same process. For example, mesophyll-specific expression of PEPC and CA in Flaveria bidentis and bundle sheath-specific expression of NAD-ME1, NAD-ME2 and mitochondrial MDH in Gynandropsis gynandra are regulated by pairs of cis-elements with high sequence homology (Gowik et al., 2004, 2017) (Reyna-Llorens et al., 2018). Moreover, PEPC and CA are co-ordinately regulated by trimethylation of histone H3K4 (Heimann et al., 2013). A comparative analysis of transcriptomes from rice and maize leaf developmental gradients predicted 118 transcription factors as candidate regulators of C_4_ gene expression (Wang et al., 2014). Amongst these, ZmMYB138 and ZmSBP6 were also predicted by our pipeline to regulate mesophyll- and bundle sheath-preferential clusters of genes respectively. Our analysis also identified three positive (ZmDOF2, ZmMADS1 and ZmDOF21) and two negative regulators (ZmBBR3 and ZmBBR4) as strong candidates for determining preferential expression of photosynthesis genes in the bundle sheath. In the analysis of rice and maize transcriptomes (Wang et al., 2014), DOF-binding cis-elements (WAAAG; W = T/A) were also enriched in bundle sheath-specific genes and it was proposed that they have been recruited from the ancestral C_3_ state to drive bundle sheath-specific expression. Different predictions from the two studies are likely explained by the nature of the transcriptomic datasets used. For example, it is possible that analysis of transcriptomes from rice and maize (Wang et al., 2014) identified regulators that establish differences between the C_3_ and C_4_ systems, whereas the sampling strategy in our case was able to predict genes that maintain and fine-tune cell-preferential gene expression over the photoperiod.

In maize the C_4_ acid decarboxylases NADP-ME and PEPCK drive malate and aspartate metabolism in bundle sheath cells as sources of CO_2_ for RuBisCO in the Calvin-Benson-Bassham cycle (Chang et al., 2012; P. Li et al., 2010; Tausta et al., 2014). Our understanding of how NADP-ME and PEPCK genes are transcriptionally regulated in C_4_ plants is limited. To date, only ZmbHLH128 and ZmbHLH129 were shown to bind the maize NADP-ME promoter in vivo (Borba et al., 2018; Schlüter & Weber, 2020). Our pipeline identified DOF2 as a candidate activator of diel and bundle sheath-preferential expression of NADP-ME, and MADS1 as an activator of PEPCK and RBCS. Transactivation assays confirmed interaction between these transcription factors and promoters of the C_4_ genes in planta. Notably, DOF2 in maize has previously been shown to repress transcription of the C_4_ PEPC gene (Yanagisawa, 2000; Yanagisawa & Sheen, 1998). Our findings therefore suggest that maize DOF2 plays a dual-function in the regulation of C_4_ genes in bundle sheath cells through repression of PEPC and activation of NADP-ME. Despite transcription factors often being classified as ‘activators’ or ‘repressors’, some can have both roles depending on the cis-regulatory element to which they bind, the structure of the surrounding chromatin, protein post-translational modifications and interaction with other proteins (Boyle & Després, 2010).

The work reported here extends our understanding of C_4_ regulation. For example, the diel and spatial patterning of RBCS in C_4_ is well-characterised and known to be controlled by multiple levels of gene regulation, including transcriptional and post-transcriptional (Berry et al., 1986; Borello et al., 1993; Giuliano et al., 1988; M. Patel et al., 2004; Minesh Patel et al., 2006; Xu et al., 2001). In maize the RBCS gene is transcriptionally regulated by two independent cis-elements present in untranslated regions (UTRs). In the 5’ UTR an I-box is essential for light-mediated activation (Giuliano et al., 1988) whilst in the 3’ UTR a HOMO motif, which binds the Transcription Repressor-Maize 1 protein, drives mesophyll-repression (Xu et al., 2001). The data presented here identify MADS1 as an additional regulatory element associated to the diel expression of RBCS. It seems likely that MADS1 activates RBCS gene expression in bundle sheath cells as both are positively co-expressed with MADS1 and RBCS peaking at six hours and ten hours after dawn respectively. Combined with previous findings, our data therefore suggest that bundle sheath-preferential expression of RBCS is achieved through HOMO-mediated repression of RBCS transcription in mesophyll (Xu et al., 2001) combined with MADS1-mediated activation of RBCS in bundle sheath.

More broadly, our findings are consistent with previous knowledge that MADS-domain transcription factors are key components of genetic regulatory networks involved in plastic developmental responses in plants (Castelán-Muñoz et al., 2019). MADS1 also enhanced expression of PEPCK and so it seems likely, that as with RBCS, PEPCK requires additional regulatory elements to allow modulation of cell-preferential gene expression and induction by light. In summary, we report that in maize the extent to which C_4_ genes are expressed in either mesophyll cells or bundle sheath strands varies during the day. The distinct dynamics of transcript abundance between the two cell-types allowed us to undertake a gene co-expression analysis that together with trans-activation assays in planta showed that DOF2 and MADS1 act as transcriptional activators of diel and bundle sheath-preferential expression of C_4_ genes. It was also noticeable that cell-preferential expression of C_4_ genes either preceded or were coincident with maximum rates of photosynthesis.

## Materials and Methods

### Growth conditions and photosynthetic measurements

Zea mays L. var. B73 plants were grown in M3 High Nutrient soil (Levington Advance) fertilised with 1 g L^-1^ Osmocote, under 16-hours light photoperiod, 26°C day and night, 55% relative humidity and ambient carbon dioxide (CO_2_) concentration. A light-emitting diode (LED) panel provided light at ∼500 μmol m^-2^ s^-1^ Photosynthetic Photon Flux Density. Fully expanded third leaves of 10-day-old maize plants were used for all analyses.

CO_2_ assimilation and chlorophyll fluorescence of fourteen 10-day-old maize third leaves were measured simultaneously with a portable gas-exchange system LI-6800 (LI-COR Biosciences) equipped with a Fluorometer head 6800-01 A (LI-COR Biosciences). Leaves were first equilibrated at 400 ppm CO_2_, an irradiance of 500 μmol m^-2^ s^-1^, red-blue actinic light (90%/10%), leaf temperature 25°C, 15 mmol mol^-1^ H_2_O, and a flow rate 500 μmol s^-1^. Effective quantum yield of Photosystem II (φPSII) was probed simultaneously with the gas-exchange measurements under red-blue actinic light (90%/10%) using a multiphase saturating flash routine (Loriaux et al., 2013) with phase 1 and 3 at 8000 μmol m^-2^ s^-1^. Maize leaves were dark-adapted for 4 hours prior to obtaining F_o_ and F_m_, the minimal and maximal levels of fluorescence, respectively.

For measurements of chlorophyll fluorescence in diel and constant light and temperature, (26°C day and night/subjective night), fragments of six 10-day-old maize third leaves were excised and placed into individual wells of a black 96-well imaging plate (Greiner) filled with of 0.8% (w/v) bactoagar, ½ MS, 0.5 µM 6-benzyl-aminopurine adjusted to pH5.7 with 0.5 M KOH and 0.5 M HCl. The plate of leaf fragments was then moved to a CFimager (Technologica Ltd) and allowed to acclimate under 100 μmol m^-2^ s^-1^ blue light until dusk when lights were switched off. At dawn of the following day a light regime was used to capture ‘day’ images which consisted of 20 minutes darkness; 800 ms saturating pulse of 6172 μmol m^-2^ s^-1^ blue light, 40 minutes blue light at irradiance 100 μmol m^-2^ s^-1^, 800 ms saturating pulse of 6172 μmol m^-2^ s^-1^ blue light, which was repeated every hour. After 16 hours the blue light source was switched off and a single 800 ms saturating pulse of 6172 μmol m^-2^ s^-1^ blue light was applied once per hour to capture “night” images. At dawn of the next day this repeating light regime was run continuously for a further 72 hours to simulate constant light but with dark breaks to allow imaging as has been used previously (Wittern et al., 2023). Chlorophyll fluorescence parameters were calculated using the image scripts provided by the manufacturer. The empirical p-values and free running period estimates associated with each parameter were calculated from linear detrended data collected between timepoints 48-96 hours in repeating light using the meta.meta function in the MetaCycle R-package (Wu et al., 2016).

### Mesophyll and bundle sheath strand isolation, RNA extraction and sequencing

Fully expanded segments of 10-day-old maize third leaves were harvested at 0, 2, 6, 10, 14, 18 and 22 hours across the photoperiod. The top 0.5 cm of each leaf was discarded, and the midrib removed. Mesophyll extracts were isolated as described previously by Covshoff, Furbank, Leegood, & Hibberd (2013) and bundle sheath strands according to Markelz, Costich, & Brutnell (2003) and John, Smith-Unna, Woodfield, Covshoff, & Hibberd (2014). Three replicates of six leaves each were initially rolled to extract mesophyll sap and then blended to isolate bundle sheath strands. Mesophyll sap was rapidly collected and deposited into RLT lysis buffer for RNA extraction (RNeasy Plant Mini Kit, Qiagen). Excess moisture was removed of the purified bundle sheath strands on a bed of paper towel. Bundle sheath strands were flash frozen in liquid nitrogen and stored at −80°C prior to RNA extraction.

Total RNA was extracted from three independent samples of mesophyll- and bundle sheath-enriched tissues collected at seven time-points (42 samples) using RNeasy Plant Mini Kit (Qiagen). To eliminate residual genomic DNA, the RNA was treated with TURBO DNA-free kit (Ambion) following the manufacturer’s instructions. Initial quality control of total RNA was performed by a photometric measurement on a NanoDrop 1000 device. This was followed by RQN determination via a Fragment Analyzer System (AATI) using the DNF-471 standard sensitivity RNA Assay. Final RNA quantification was performed by a fluorometric Qubit assay (RNA HS, ThermoFisher Scientific). Library preparation was carried out on a PerkinElmer Sciclone NGS robotics unit using the Illumina TruSeq stranded mRNA sample Preparation Kit (#15031047 Rev.E) following the manufacturer’s instructions. Input amount of total RNA was 200 ng. Final libraries were passed through an additional bead clean-up step in a 1:1 ratio (sample/beads) to remove primer dimers. Quality control on a Fragment Analyzer System (AATI) was used to determine fragment length distribution using the DNF-474 Assay. For quantification purposes, a fluorometric Qubit dsDNA HS Assay Kit was used. Libraries were diluted to 2 nM prior to equimolar pooling into 6 separate pools which were then each sequenced on individual flow cell lanes. Paired-End sequencing with a 2×150 bp read length was performed on an Illumina HiSeq3000 system using the HiSeq 3000/4000 PE Cluster Kit (PE-410-1001) and the HiSeq 3000/4000 SBS Kit 300 cycles (FC-410-1003). Clustering and sequencing were carried out following to the manufacturer’s instructions. Library preparation and sequencing were done at the Genomics and Transcriptomics Labor of the University of Düsseldorf.

### Read assembly, annotation, and quantification of transcript abundance

Reads were mapped to the Zea mays B73 genome AGPv3 (from Ensembl Plants, http://plants.ensembl.org) and quantified as Transcripts per Million (TPM) (Wagner et al., 2012) using RSEM version 1.2.23 with default settings (B. Li & Dewey, 2011) in conjunction with Bowtie 1 (Langmead et al., 2009). Differential expression analysis was performed using the DESeq2 R package (Love et al., 2014) with read counts used as input. Cell-type was treated as condition (mesophyll vs. bundle sheath). Benjamini-Hochberg corrected p-value was set to < 0.01 to identify differentially expressed genes (Supplemental Table 2) (Benjamini & Hochberg, 1995).

### Data analysis and visualisation

Data analysis was performed using R (R Development Core Team, 2009) unless stated otherwise. The R package ggplot2 (Wickham, 2009) was used to generate all graphs. Principal Component Analysis was performed on the mean of transcriptome triplicates of mesophyll and bundle sheath samples collected at 0, 2, 6, 10, 14, 18, and 22 hours. The Pearson’s correlation coefficient was calculated between transcriptomes of three biological replicates from mesophyll and bundle sheath samples. K-means clustering was performed on expressed genes (TPM > 5). Genes were quantile normalized and transformed to Z-score values. A total of fifteen centres were selected based on the total within sum of squares. Gene Ontology (GO) term enrichment analysis was performed using AgriGO v2 [GO analysis toolkit and database for agricultural community (Tian et al., 2017)] with the following settings: statistical test method – Fisher; Multi-test adjustment method – Hochberg (FDR); gene ontology type – Complete GO. A False Discovery Rate cutoff of ≤ 0.01 was set to identify significantly enriched GO terms in clusters of co-expressed genes (detailed in Supplemental Table 4).

Genes encoding maize transcription factors were downloaded from PlantTFDB v4.0 [2331 genes, http://planttfdb.cbi.pku.edu.cn, (Jin et al., 2017)] and Grassius [2605 genes, http://www.grassius.org, (Yilmaz et al., 2009)]. Only the 2110 genes present in both databases were considered in further analyses. Maize genes encoding transcription factors were assigned into families according to PlantTFDB v4.0. Motif enrichment analysis across genes was performed for each cluster using the “Analysis of Motif Enrichment” tool from the MEME suite (Bailey et al., 2009; McLeay & Bailey, 2010) using default parameters. For each transcript present in a particular cluster, promoter sequences (−2kb to +0.5kb from the transcription start site) were retrieved and used as input. Control sequences were defined as the entire set of sequences (all clusters) minus those sequences present in the cluster of interest. Gene co-expression network was built using Cytoscape (Shannon et al., 2003).

Maize orthologs were identified for circadian clock genes from Arabidopsis thaliana using OrthoFinder (Emms & Kelly, 2019) including the proteomes of seven representative plant species (Arabidopsis thaliana, Oryza sativa, Triticum aestivum, Brachypodium distachyon, Setaria italica, Sorghum bicolor and Zea mays). Proteomes were downloaded from the ENSEMBL website (www.ensembl.com). Phylogenetic trees were generated using Dendroscope (www.dendroscope.org; Huson, Rupp, Berry, Gambette, & Paul, 2009).

### Trans-activation assays in planta

Constructs were generated using Golden Gate cloning as described in Supplemental Table 8. Coordinates of the NADP-ME (GRMZM2G085019) and PEPCK (GRMZM2G001696) promoters (1.5 Kb upstream of the translation start site) enriched in DOF2 and MADS1 motifs, respectively, were retrieved from the motif enrichment analysis. For the trans-activation assays with the NADP-ME promoter, two fragments of 106 bp that contain a 6 bp-DOF2 motif (‘aaagcc’ in NADP-MEa and ‘ggcttt’ in NADP-MEb) flanked by 50 bp-endogenous promoter sequence either side of the motif were cloned upstream of a minimal 35S promoter (Supplemental Table 8). For the PEPCK promoter, one fragment of 121 bp that contain a 21 bp-MADS1 motif (‘tttctttcttttgttctccgc’) flanked by 50 bp-endogenous promoter sequence either side of the motif was cloned upstream of a minimal 35S promoter (Supplemental Table 8). For the RBCS promoter, one fragment of 121 bp that contained a 21 bp-MADS1 motif (‘aaacgaaaaaaataacaaaca’) flanked by 50 bp-endogenous promoter sequence either side of the motif was cloned upstream of a minimal 35S promoter (Supplemental Table 8). 106 bp-<u>pNADP-MEa</u>: −586 to −692 bp upstream of the translation start site with the 6 bp-DOF2 motif (‘aaagcc’) at −636 to −642 bp upstream of the translation start site; 106 bp-<u>pNADP-MEb</u>: −112 to −218 bp upstream of the translation start site with the 6 bp-DOF2 motif (‘ggcttt’) at −162 to −168 bp upstream of the translation start site; 121 bp-<u>pPEPCK</u>: −815 to −936 bp upstream of the translation start site with the 21 bp-MADS1 motif (‘tttctttcttttgttctccgc’) at −865 to −886 bp upstream of the translation start site; 121 bp-<u>pRBCS</u>: −1089 to −968 bp upstream of the translation start site with the 21bp-MADS1 motif (‘aaacgaaaaaaataacaaaca’) at −1039 to −1018 bp upstream of the translation start site. Level 1 constructs were made such that these promoter fragments were placed upstream of the GUS reporter gene. To produce level 2 constructs, these were combined with a transformation control containing the LUCIFERASE reporter driven by the constitutive NOS promoter, the transcription factor of interest driven by the constitutive LjUBI promoter and the P19 silencing suppressor under control of the CaMV35S promoter. Constructs were transformed into the Agrobacterium tumefaciens strain GV3101. Overnight cultures of A. tumefaciens were pelleted and resuspended in infiltration buffer [10 mM MES (pH 5.6), 10 mM MgCl_2_, 150 μM acetosyringone] to an optical density of 0.3. Cultures were then incubated for 2 hours at room temperature and infiltrated into the abaxial side of leaves of four-week-old Nicotiana benthamiana plants with 1 mL syringe. Leaf discs from the infiltrated regions were sampled 48 hours after infiltration and flash frozen in liquid nitrogen. Protein for the 4-methylumbelliferyl-b-D-glucuronide (MUG) and luciferase (LUC) assays was extracted in 1x passive lysis buffer (PLB: Promega). MUG assays were performed by adding 40 mL of protein extract to 100 mL of MUG assay buffer [2 mM MUG, 50 mM NaH_2_PO_4_ /Na_2_ HPO_4_ buffer (pH 7.0), 10 mM EDTA, 0.1% (v/v) Triton X-1000, 0.1%(w/v) sodium lauroyl sarcosinate, 10 mM DTT]. Stop buffer (200 mM Na_2_CO_3_) was added at 0 and 120 minutes, and the rate of MUG accumulation was measured in triplicate on a plate reader (CLARIOstar, BMG lab tech) with excitation at 360 nm and emission at 465 nm. LUC activity was measured with 20 mL of protein sample and 100 mL of LUC assay reagent (Promega). Promoter activation was calculated as (rate of MUG accumulation / LUC luminescence) x 100.

### Accession numbers

All referenced gene names and accessions are detailed in Supplemental Tables 2, 3, 4 and 6. RNA-sequencing data generated in this study have been deposited to the National Center for Biotechnology Information Sequence Read Archive with accession number PRJNA635519.

## Acknowledgements

We thank André M. Cordeiro, Bruno Alexandre and Joana Rodrigues (ITQB-NOVA, Oeiras, Portugal) for helping in mesophyll and bundle sheath cell isolation. This work was supported by the European Union project 3to4 (Grant agreement no: 289582). By Fundação para a Ciência e Tecnologia (FCT) through research unit GREEN-it ‘Bioresources for Sustainability’ (UID/Multi/04551/2013, UIDB/04551/2020, UIDP/04551/2020). A.R.B. (SFRH/BD/105739/2014), A.M.G. (SFRH/BD/89743/2012) and N.J.M.S. (IF/01126/2012 – POPH-QREN). By ERACAPS grant C4BREED and BBSRC grants BB/L014130, BBP0031171 and BB/S006370/1. By ERC Grant Revolution RG80867. For the purpose of open access, the authors have applied a Creative Commons Attribution (CC BY) licence to any Author Accepted Manuscript version arising from this submission.

A.R.B., I.R-L., J. K., A.A.R.W., N.J.M.S. and J.M.H. conceptualised the experiments. A.R.B., I.R-L. and G.S. performed photosynthetic measurements. A.R.B., P.G., A.G. and N.J.M.S isolated mesophyll and bundle sheath cells. A.R.B., I.R-L. and P.J.D. conducted data analysis. A.R.B. and P.J.D. performed trans-activation assays in planta. A.R.B., I.R-L. and J.M.H. wrote the article and prepared the figures.

## Figure Legends

**Supplemental Figure 1.** Heatmap of Pearson’s correlation coefficient (PCC) calculated between transcriptomes of biological replicates 1, 2 and 3 of mesophyll and bundle sheath samples collected at time-points 0, 6, 10, 14, 18 and 22 hrs. High to low values of Pearson’s correlation coefficient are shown as red to blue.

**Supplemental Figure 2.** Components of the maize circadian oscillator. A) Species tree inferred by Orthofinder with bootstrap values displayed at each node. B-C) Orthologue trees inferred for PRR7 (B) and PRR3/5/9 (C). D) Diel transcript abundance profile of genes for the circadian oscillator in mesophyll and bundle sheath cells. x-axis represents time-points and y-axis transcript abundance. TPM represents Transcripts Per Million reads. White and black bars on x-axis denote light and dark periods, respectively. CCA1/LHY.1, GRMZM2G014902; CCA1/LHY.2, GRMZM2G474769; PRR7.1, GRMZM2G005732; PRR7.2, GRMZM2G033962; PRR7.3, GRMZM2G095727; PRR3/5/9.1, GRMZM2G179024; PRR3/5/9.2, GRMZM2G367834; ELF3, GRMZM2G045275; LUX, GRMZM2G067702; TOC1.1, GRMZM2G148453; TOC1.2, GRMZM2G020081; TOC1.3, GRMZM2G066638; TOC1.4, GRMZM2G145058; TOC1.5, GRMZM2G174083; TOC1.6, GRMZM2G365688.

**Supplemental Figure 3.** Photosynthetic parameters measured for maize leaf fragments under one light-dark cycle (16 hrs light: 8 hrs dark) followed by 72 hours of a light regime that consisted of cycles of 40 minutes light and 20 minutes darkness. Data shown as mean with standard error (n = 6 biological replicates). A) F_m_: maximum possible yield of fluorescence, B) Linear detrended F_v_/F_m_: maximum quantum efficiency of Photosystem II (PSII) photochemistry, C) φPSII: operating efficiency of PSII, and D) F_v_’/F_m_’: maximum efficiency of PSII photochemistry in the light. Black and grey bars represent dark period and subjective night, respectively. Empirical p-values calculated using the meta.meta function in MetaCycle from timepoints 48-96 hours in repeating light where emp p < 0.01 is considered rhythmic.

**Supplemental Figure 4.** Distribution of biological processes across the diel time-course and between mesophyll and bundle sheath cells. Dot plot showing the categories of biological processes with highest significance for each cluster (FDR ≤ 0.01). Clusters 1 to 3 peaked from dawn to 2 hours of light, clusters 5 to 8 from 6 to 10 hrs, clusters 9 to 13 from 14 to 22 hrs, cluster 14 at dawn and 22 hrs, and cluster 15 from dawn to 22 hrs. Gene ratio represents the proportion of genes assigned to a functional category in a cluster. M and BS represent mesophyll and bundle sheath cells, respectively.

**Supplemental Figure 5.** Line plots representing the diel transcript abundance profile of genes encoding the cognate transcription factors for DNA-binding motifs enriched in mesophyll- and bundle sheath-preferential clusters of co-expressed genes. The x-axis represents time-points and the y-axis TPM values. TPM represents Transcripts Per Million reads. White and black bars in the x-axis denote light and dark periods, respectively. M and BS represent mesophyll and bundle sheath cells, respectively. CPP-transcription factor 1 (CPP1), GRMZM2G153754; G2-like-transcription factor 56 (GLK56), GRMZM2G067702; MYB-transcription factor 138 (MYB138), GRMZM2G139688; DNA-binding One Zinc Finger 21 (DOF21), GRMZM2G162749; MYB-transcription factor 14 (MYB14), GRMZM2G172327; HSF-transcription factor 19 (HSFTF19), AC216247.3_FG001; SBP-transcription factor 6 (SBP6), GRMZM2G138421; KNOTTED 1 (KN1), GRMZM2G017087; ABI3-VP1-transcription factor 19 (ABI19), GRMZM2G035701; NLP-transcription factor 13 (NLP13), GRMZM2G053298; SBP-transcription factor 17 (SBP17), GRMZM2G156756; HSF-transcription factor 8 (HSFTF8), GRMZM2G164909; HSF-transcription factor 4 (HSFTF4), GRMZM2G125969; BBR/BCP-transcription factor 3 (BBR3), GRMZM2G164735; BBR/BCP-transcription factor 4 (BBR4), GRMZM2G118690; Dwarf Plant 8 (D8), GRMZM2G144744; MADS-domain protein 1 (MADS1), GRMZM2G171365; DNA-binding One Zinc Finger 2 (DOF2), GRMZM2G009406; Viviparous 1 (VP1), GRMZM2G133398.

**Supplemental Figure 6.** Gene co-expression network built from RNA-seq data and DNA motif enrichment analysis. A) Transcripts encoding transcription factors (TF) with DNA-binding motif hits in photosynthesis (PS) genes (C_4_ genes and Calvin-Benson-Bassham cycle genes) were filtered by their expression levels [Transcripts Per Million reads (TPM) > 5] and a gene co-expression network built for TF and PS genes using Pearson’s correlation coefficient (cutoffs of < 0.3 and > −0.3). B) Gene co-expression network for TF and bundle sheath-preferential PS genes in clusters 7 and 3. Nodes represent TF (grey) and PS genes present in clusters 7 (dark blue) and 3 (light blue). Edges represent positive (green) and negative (red) co-expression based on the Pearson’s correlation coefficient (PCC).

