## Supplementary material for "Compartmentation of photosynthesis gene expression between mesophyll and bundle sheath cells of C_4_ maize is dependent on time of day": S Figures

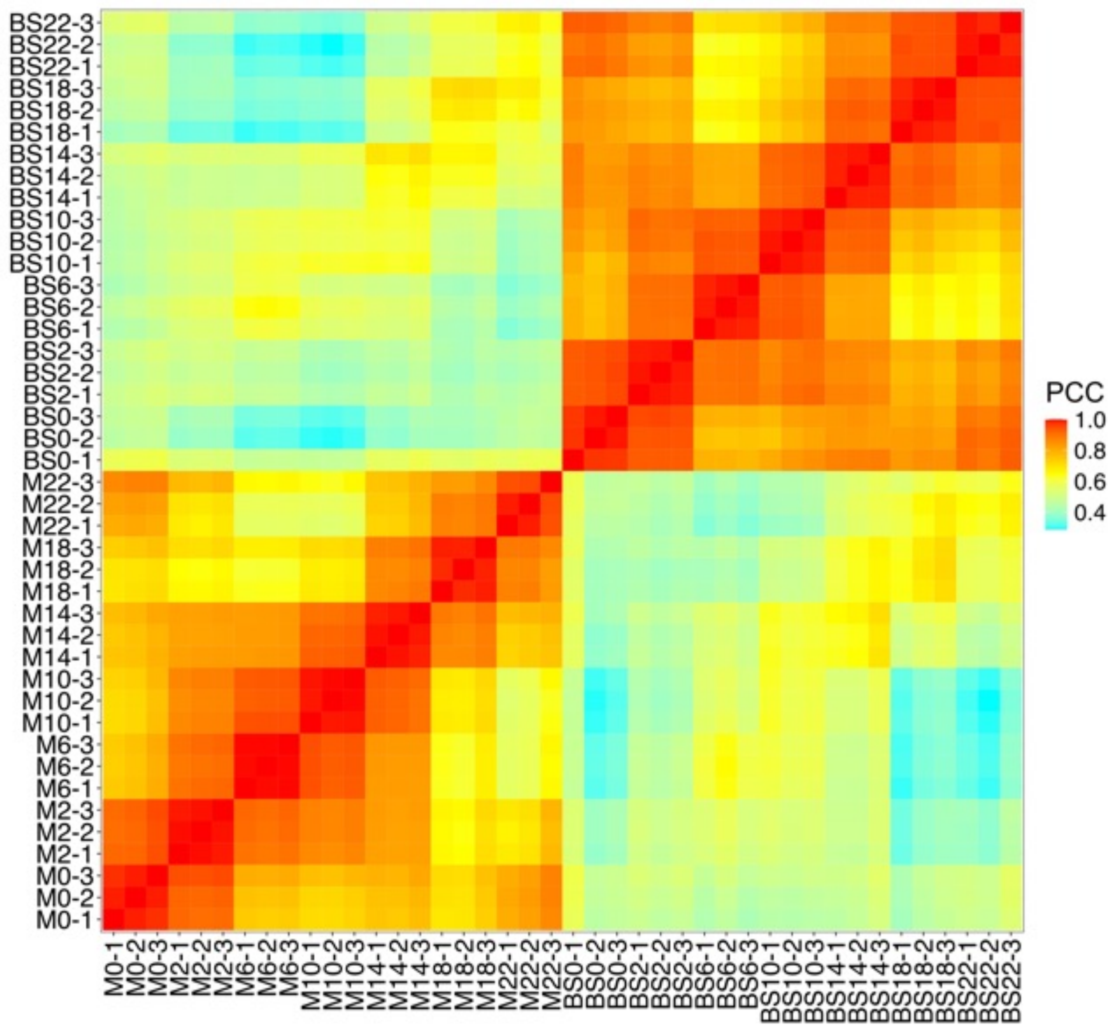


**Supplemental Figure 1.** Heatmap of Pearson's correlation coefficient (PCC) calculated between transcriptomes of biological replicates 1, 2 and 3 of mesophyll and bundle sheath samples collected at time-points 0, 6, 10, 14, 18 and 22 hrs. High to low values of Pearson's correlation coefficient are shown as red to blue.

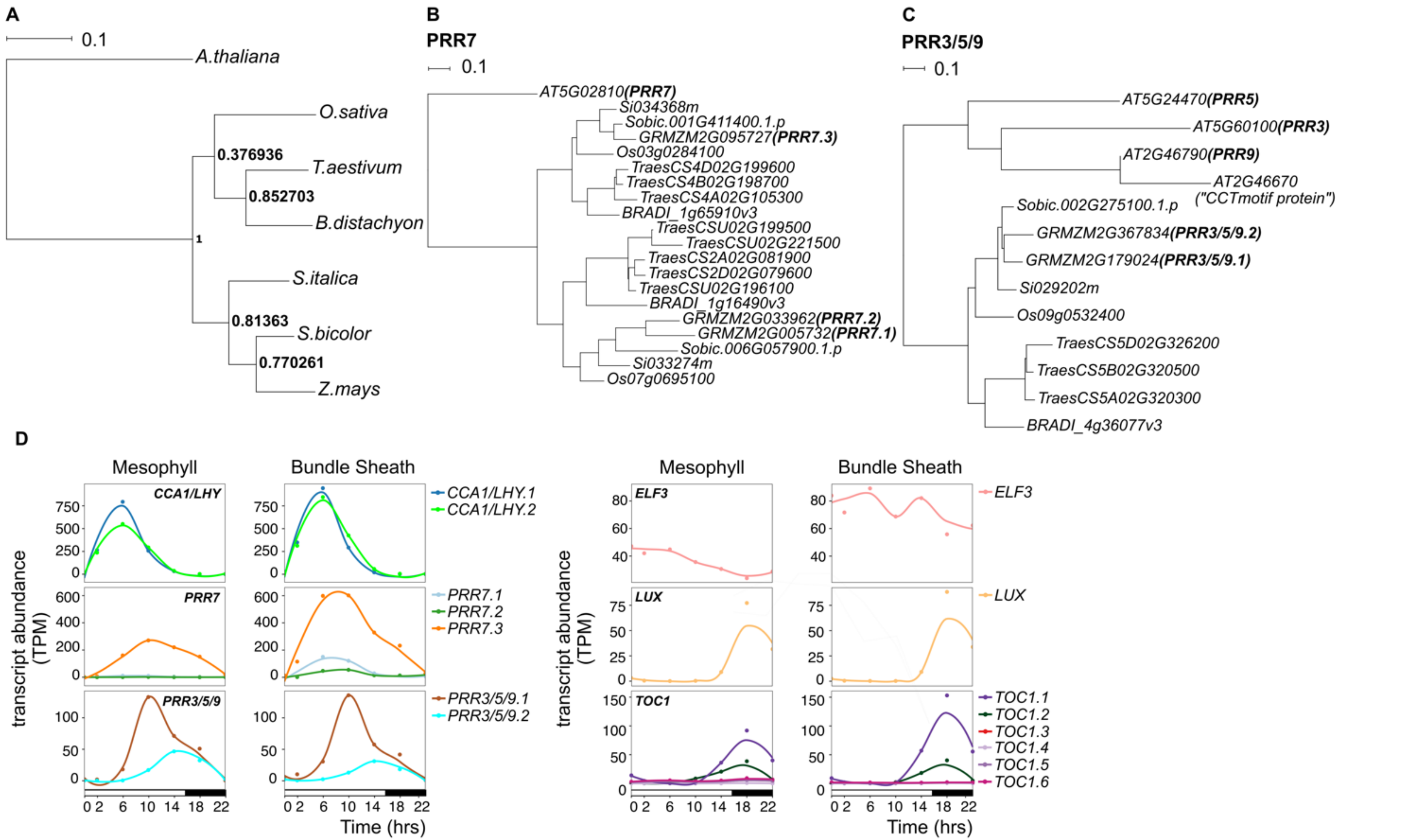


**Supplemental Figure 2**. Components of the maize circadian clock. **A)** Species tree inferred by *Orthofinder* with bootstrap values displayed at each node. **B-C)** Orthologue trees inferred for *PRR7* (B) and *PRR3/5/9* (C). **D)** Diel transcript abundance profile of core clock genes in mesophyll and bundle sheath cells. *x*-axis represents time-points and *y*-axis transcript abundance. TPM represents Transcripts Per Million reads. White and black bars on *x*-axis denote light and dark periods, respectively. *CCA1/LHY.1*, GRMZM2G014902; *CCA1/LHY.2*, GRMZM2G474769; *PRR7.1*, GRMZM2G005732; *PRR7.2*, GRMZM2G033962; *PRR7.3*, GRMZM2G095727; *PRR3/5/9.1*, GRMZM2G179024; *PRR3/5/9.2*, GRMZM2G367834; *ELF3*, GRMZM2G045275; *LUX*, GRMZM2G067702; *TOC1.1*, GRMZM2G148453; *TOC1.2*, GRMZM2G020081; *TOC1.3*, GRMZM2G066638; *TOC1.4*, GRMZM2G145058; *TOC1.5*, GRMZM2G174083; *TOC1.6*, GRMZM2G365688.

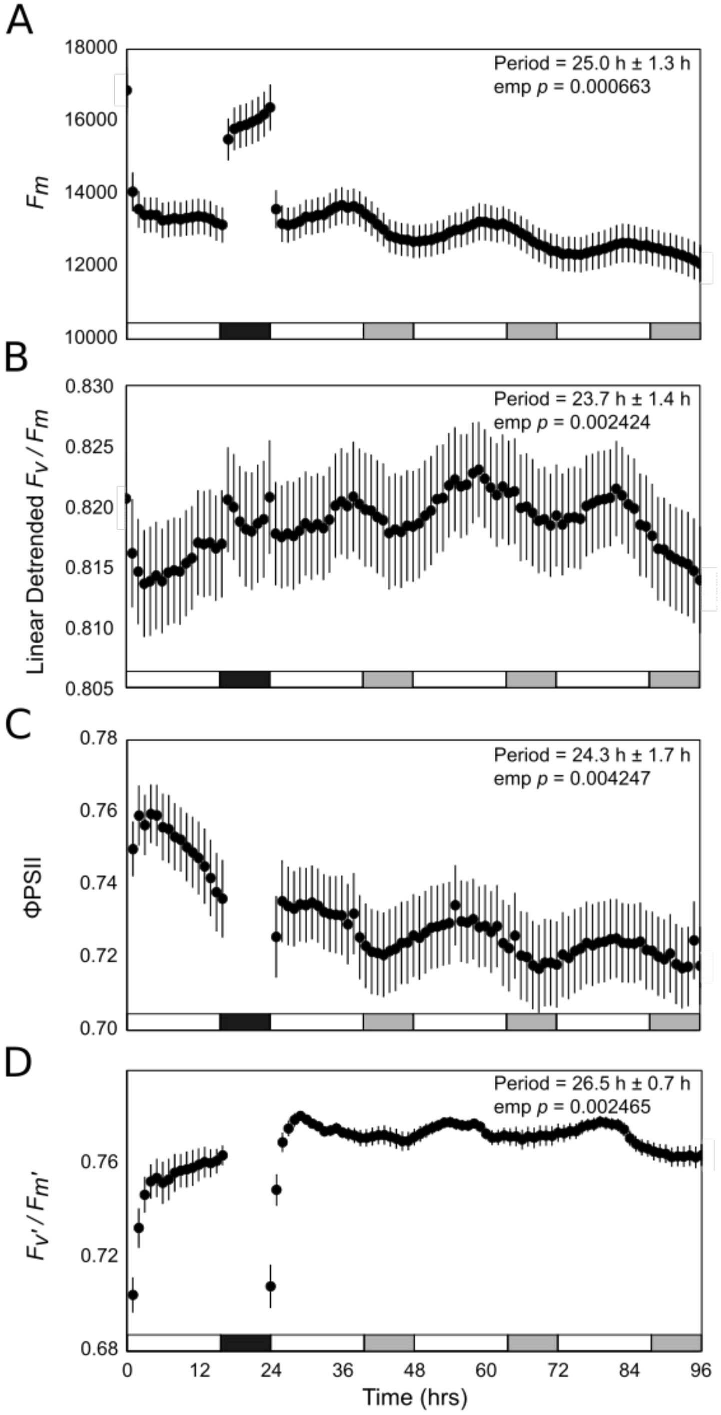


**Supplemental Figure 3.** Photosynthetic parameters measured for maize leaf fragments under one light-dark cycle (16 hrs light : 8 hrs dark) followed by 72 hours of a light regime that consisted of cycles of 40 minutes light and 20 minutes darkness. Data shown as mean with standard error (*n* = 6 biological replicates). **A)** $F_m$: maximum possible yield of fluorescence, **B)** Linear detrended $F_v/F_m$: maximum quantum efficiency of Photosystem II (PSII) photochemistry, **C)** ØPSII: operating efficiency of PSII, and D) $F_v'/F_m'$: maximum efficiency of PSII photochemistry in the light. Black and grey bars represent dark period and subjective night, respectively. Empirical *p*-values calculated using the meta.meta function in MetaCycle from timepoints 48-96 hours in repeating light where emp $p < 0.01$ is considered rhythmic.

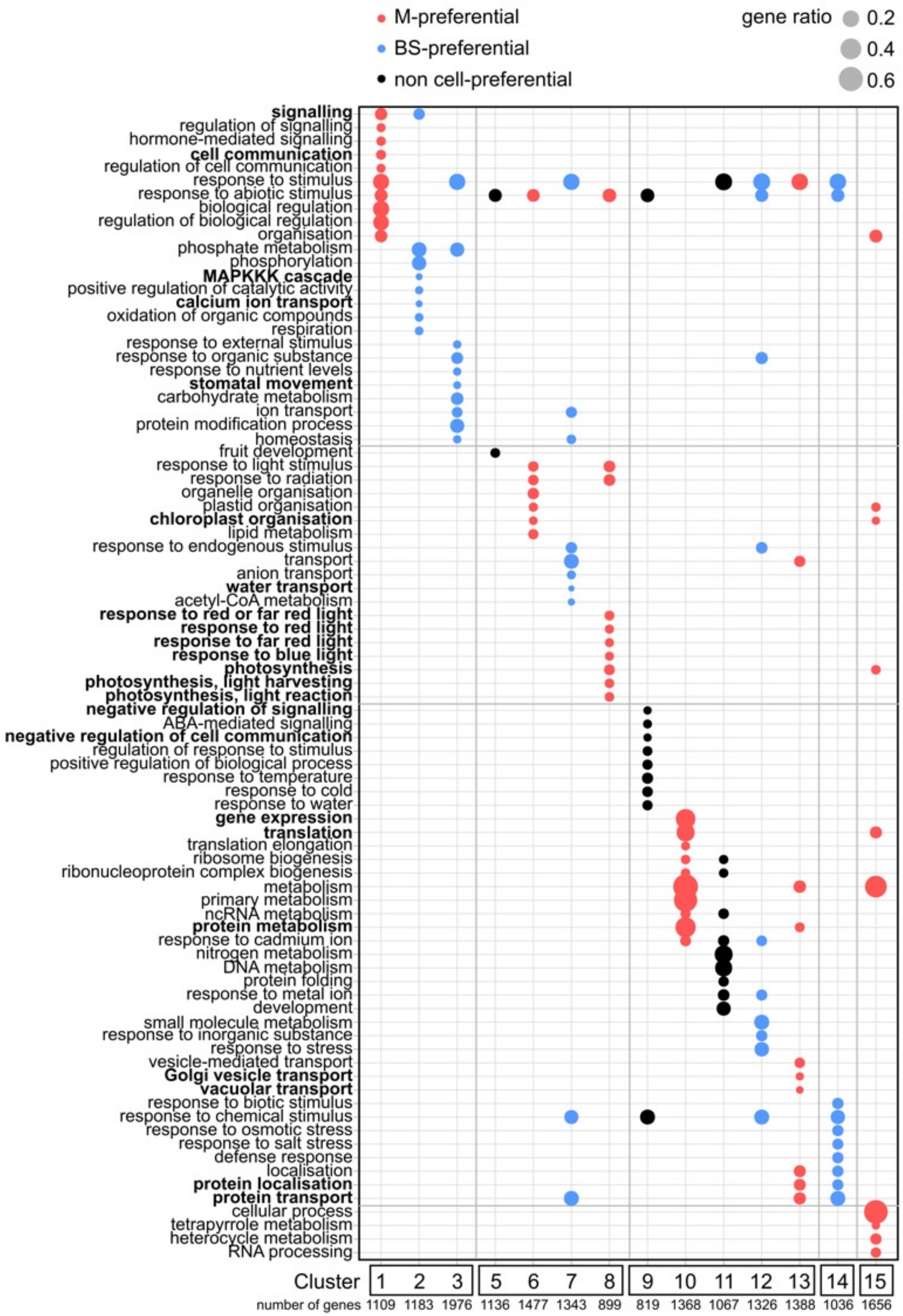


**Supplemental Figure 4.** Distribution of biological processes across the diel time-course and between mesophyll and bundle sheath cells. Dot plot showing the categories of biological processes with highest significance for each cluster (FDR ≤ 0.01). Clusters 1 to 3 peaked from dawn to 2 hours of light, clusters 5 to 8 from 6 to 10 hrs, clusters 9 to 13 from 14 to 22 hrs, cluster 14 at dawn and 22 hrs, and cluster 15 from dawn to 22 hrs. Gene ratio represents the proportion of genes assigned to a functional category in a cluster. M and BS represent mesophyll and bundle sheath cells, respectively.

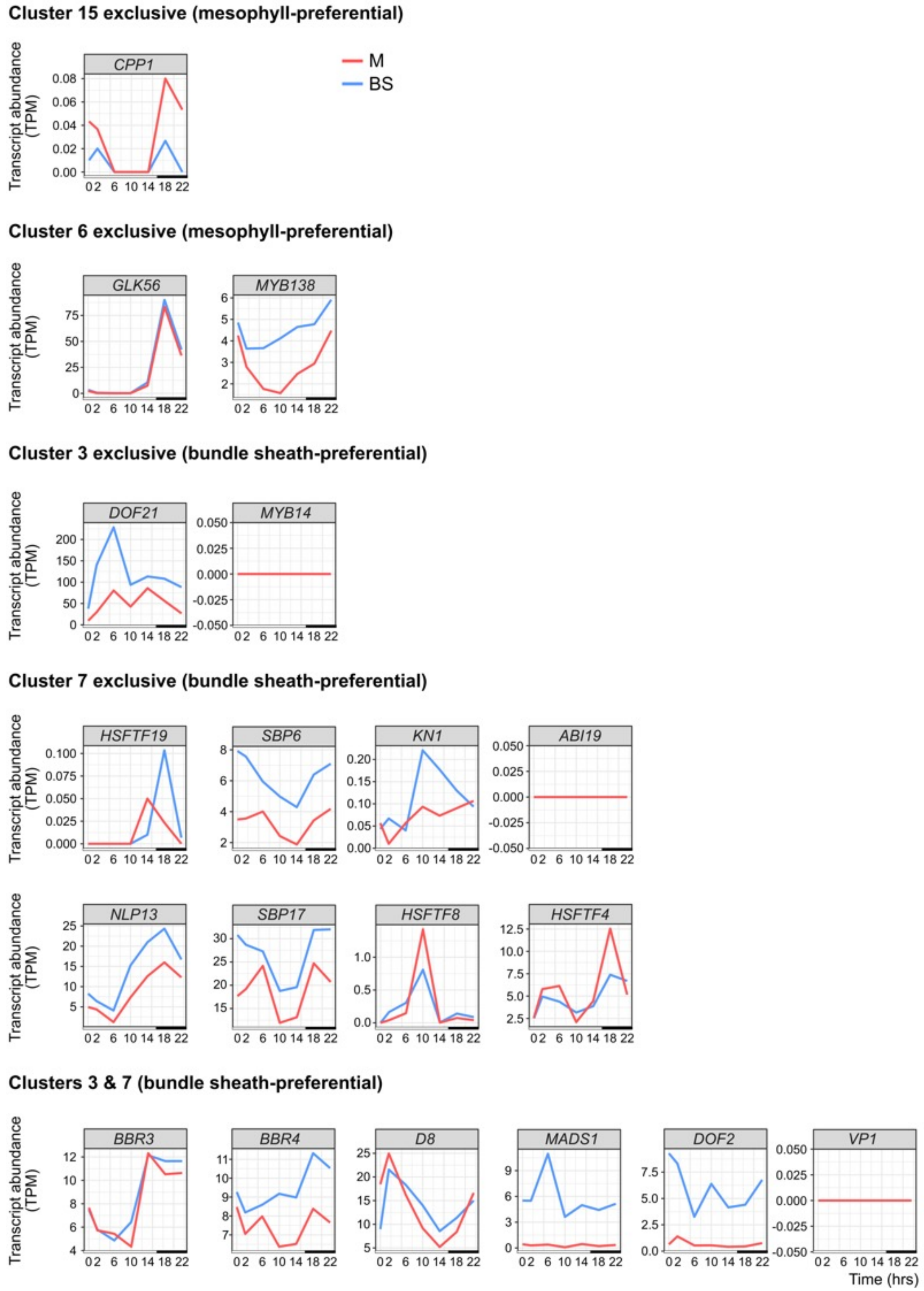


**Supplemental Figure 5**. Line plots representing the diel transcript abundance profile of genes encoding the cognate transcription factors for DNA-binding motifs enriched in mesophyll- and bundle sheath-preferential clusters of co-expressed genes. The *x*-axis represents time-points and the *y*-axis TPM values. TPM represents Transcripts Per Million reads. White and black bars in the *x*-axis denote light and dark periods, respectively. M and BS represent mesophyll and bundle sheath cells, respectively. CPP-transcription factor 1 (CPP1), GRMZM2G153754; G2-like-transcription factor 56 (GLK56), GRMZM2G067702; MYB-transcription factor 138 (MYB138), GRMZM2G139688; DNA-binding One Zinc Finger 21 (DOF21), GRMZM2G162749; MYB-transcription factor 14 (MYB14), GRMZM2G172327; HSF-transcription factor 19 (HSFTF19), AC216247.3_FG001; SBP-transcription factor 6 (SBP6), GRMZM2G138421; KNOTTED 1 (KN1), GRMZM2G017087; ABI3-VP1-transcription factor 19 (ABI19), GRMZM2G035701; NLP-transcription factor 13 (NLP13), GRMZM2G053298; SBP-transcription factor 17 (SBP17), GRMZM2G156756; HSF-transcription factor 8 (HSFTF8), GRMZM2G164909; HSF-transcription factor 4 (HSFTF4), GRMZM2G125969; BBR/BCP-transcription factor 3 (BBR3), GRMZM2G164735; BBR/BCP-transcription factor 4 (BBR4), GRMZM2G118690; Dwarf Plant 8 (D8), GRMZM2G144744; MADS-domain protein 1 (MADS1), GRMZM2G171365; DNA-binding One Zinc Finger 2 (DOF2), GRMZM2G009406; Viviparous 1 (VP1), GRMZM2G133398.

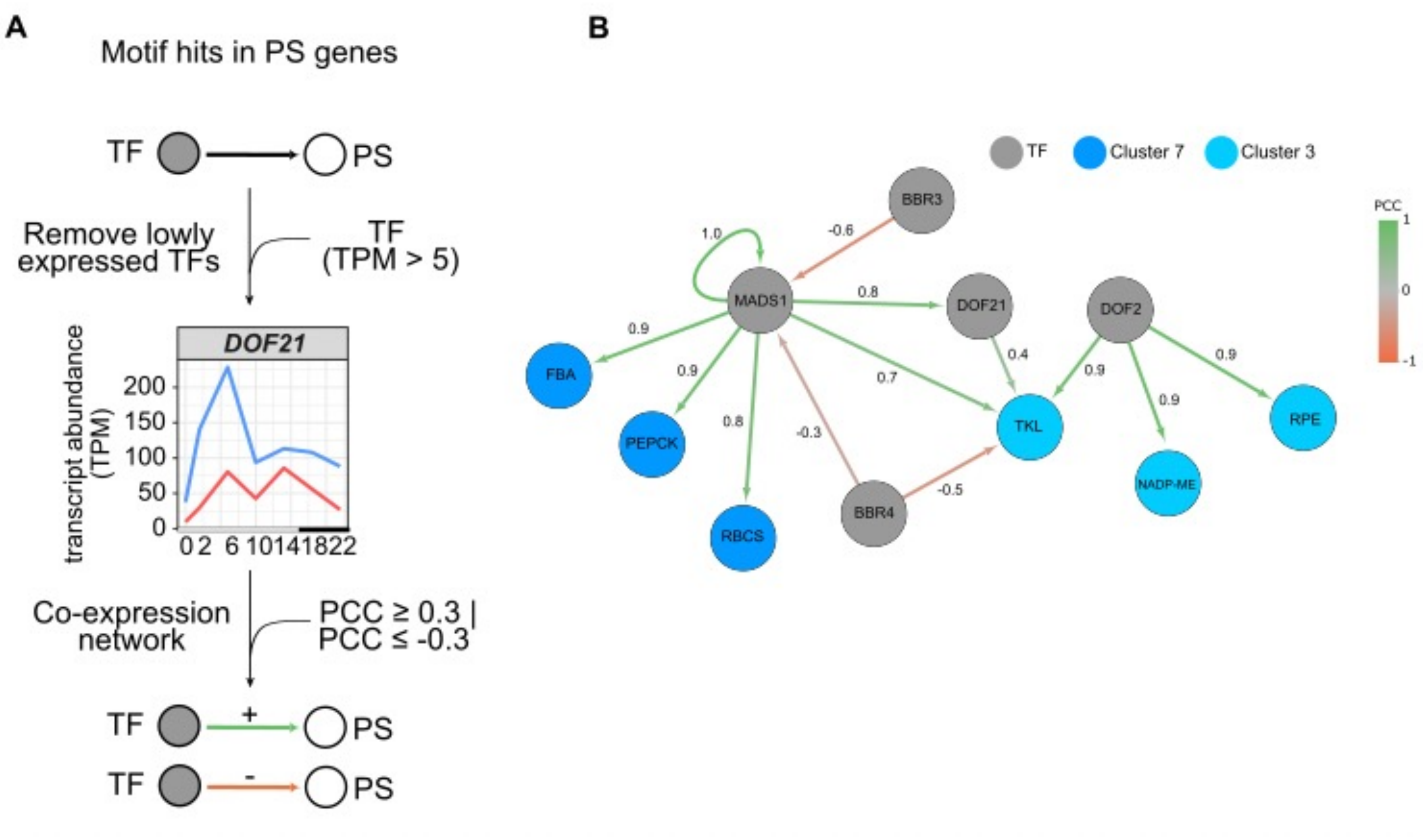


**Supplemental Figure 6.** Gene co-expression network built from RNA-seq data and DNA motif enrichment analysis. **A)** Transcripts encoding transcription factors (TF) with DNA-binding motif hits in photosynthesis (PS) genes ($C_4$ genes and Calvin-Benson-Bassham cycle genes) were filtered by their expression levels [Transcripts Per Million reads (TPM) > 5] and a gene co-expression network built for TF and PS genes using Pearson's correlation coefficient (cutoffs of < 0.3 and > -0.3). **B)** Gene co-expression network for TF and bundle sheath-preferential PS genes in clusters 7 and 3. Nodes represent TF (grey) and PS genes present in clusters 7 (dark blue) and 3 (light blue). Edges represent positive (green) and negative (red) co-expression based on the Pearson's correlation coefficient (PCC).
